# Dietary inclusion of papaya seed powder significantly modulates reproductive function and growth dynamics in Nile tilapia cultured under controlled conditions

**DOI:** 10.1101/2025.11.23.690043

**Authors:** Roba Tashome Denagde, Mimi Nuguse

## Abstract

Nile tilapia (*Oreochromis niloticus*) is a key species in aquaculture but faces challenges related to premature sexual maturity and prolific breeding. This study investigated the effects of dietary papaya seed powder (PSP) on growth performance, reproductive parameters, and survival of Nile tilapia under controlled laboratory conditions. Juvenile fish were fed diets containing 0 (control), 2, 4, and 6 g PSP/kg feed. Results demonstrated that moderate PSP inclusion (4 g/kg, T2) significantly enhanced growth, feed conversion efficiency, and survival rate (90%), whereas higher inclusion levels (6 g/kg) reduced growth. Gonad weight and gonadosomatic index (GSI) were lowest in T2, indicating potential antifertility effects, particularly in males. Water quality remained stable across all treatments, suggesting that the observed effects were primarily due to dietary PSP. These findings indicate that PSP can serve as a natural, cost-effective feed additive that promotes growth at optimal levels while suppressing reproduction at higher doses, offering a sustainable strategy for tilapia aquaculture.

## 1. Introduction

Aquaculture, the controlled cultivation of aquatic organisms such as fish, shellfish, and algae, is a critical contributor to global food security, providing affordable, high-quality protein, conserving wild fish stocks, and supporting rural livelihoods [1-3]. Among freshwater species, tilapia is one of the most widely cultured and economically significant groups. Its rapid growth, adaptability, and resilience have made Nile tilapia (*Oreochromis niloticus*) a dominant species driving the expansion of freshwater aquaculture worldwide [4, 5].

Despite these advantages, tilapia aquaculture faces challenges due to uncontrolled reproduction. Early sexual maturation and frequent breeding often result in overpopulation, stunted growth, reduced feed conversion efficiency, and ecological imbalances within production systems [6]. To address this, several reproductive control strategies including hormonal sex reversal, hybridization, manual sex separation, high-density culture, and temperature-induced monosex production have been employed [7]. However, these methods are limited by labor intensity, ethical concerns, incomplete sex reversal, and potential health and environmental risks associated with synthetic hormones.

Given these constraints, plant-based bioactive compounds have emerged as sustainable and cost-effective alternatives for reproductive regulation. *Carica papaya* L., commonly known as papaya, is a tropical fruit available year-round and rich in bioactive constituents such as papain, chymopapain, carpine, myrosin, carpasemine, alkaloids, benzyl isothiocyanate, and fatty acids including oleic, palmitic, stearic, and linoleic acids, many of which possess antifertility, antimicrobial, and antioxidant properties [8, 9].

Recent studies have highlighted the antifertility potential of papaya seed powder (PSP) in Nile tilapia. Dietary inclusion of PSP significantly reduced gonadosomatic index (GSI) and spawning frequency [10, 11], while many reported suppression of gonadal development and reproductive activity under controlled conditions. These effects are primarily attributed to bioactive compounds such as benzyl isothiocyanate and alkaloids, which interfere with steroid genesis and spermatogenesis. Concurrently, enzymes such as papain enhance protein digestibility and nutrient assimilation, potentially improving growth performance and feed utilization [12].

Although PSP demonstrates a dual role in promoting growth and regulating reproduction, optimal dietary inclusion levels remain unclear. Excessive supplementation may introduce anti nutritional factors such as tannins and saponins, which can reduce feed palatability and nutrient absorption. Therefore, determining a safe and effective PSP concentration is crucial for its sustainable application in tilapia aquaculture.

This study aimed to evaluate the effects of dietary PSP supplementation at varying levels (0, 2, 4, and 6g/kg feed) on growth performance, feed utilization, reproductive parameters, and survival of Nile tilapia under controlled indoor conditions. Water quality parameters were closely monitored to ensure that observed effects were attributable to dietary treatment rather than environmental variation. The findings are expected to provide valuable insights into the antifertility potential of papaya seed powder and its application as a natural, cost-effective, and eco-friendly feed additive for sustainable tilapia aquaculture.

## 2. Materials and Methods

### 2.1. Experimental Materials, Site, and Design

Ripe papaya (*Carica papaya* L.) fruits were procured from the local market in Mattu Town, Ethiopia. Wheat and barley brans were collected as milling by-products from the same locality. Fish meal, used as a protein source, was obtained from the Fish Production and Marketing Enterprise located at the Gilgel Gibe Reservoir.

The experimental study was conducted at the Zoological Sciences Laboratory, Department of Biology, and Mattu University, Ethiopia. A flow-through aquarium system was employed following the design described by Giri, S., et al. [13]. Eight 40 Litter glass jars were used, representing three treatment groups and one control, each with two replicates as shown in (Figure 1).

**Figure 1.**
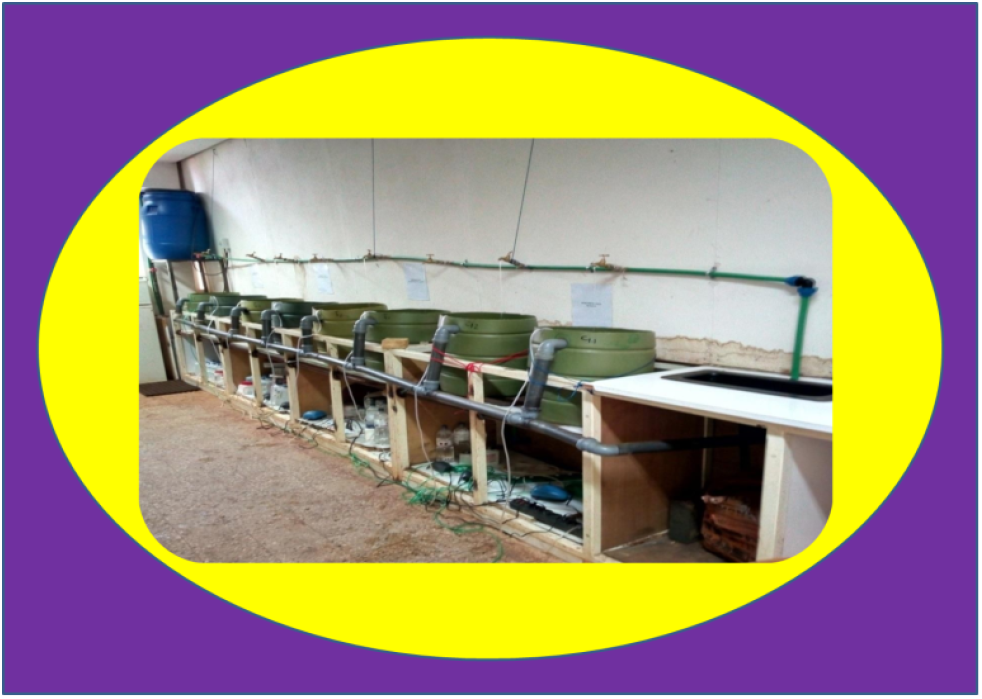
Schematic representation of the experimental setup.

A total of 80 Nile tilapia (*Oreochromis niloticus*) fingerlings (mean initial body weight: 20.11 ± 4.77 to 20.67 ± 6.39 g; mean total length: 9.96 ± 0.81 to 10.52 ± 1.01 cm) were collected from the Gilgel Gibe Reservoir and transported in partially filled plastic containers to the laboratory. The fish were acclimatized for 17 days in a glass aquarium (1.20 × 0.60 m) by gradually mixing reservoir water with dechlorinated tap water to minimize stress and ensure water-quality compatibility. During acclimatization, fingerlings were fed a basal diet at 10% of their body weight twice daily.

After acclimatization, undifferentiated fingerlings were randomly distributed at a stocking density of 10 fish per jar. Water quality was maintained through daily cleaning by siphoning out waste and feed residues, followed by replacing 50% of the water with fresh tap water. The experimental diets were offered at 10% of body weight, twice daily at 09:00 and 15:00 hours, throughout the study period [14].

The feeding trial consisted of four dietary treatments, each replicated twice. The control group (C) received a basal diet without papaya seed powder (0 g PSP/kg diet). Treatment 1 (T1) was supplemented with 2 g PSP/kg diet, Treatment 2 (T2) with 4 g PSP/kg diet, and Treatment 3 (T3) with 6 g PSP/kg diet. The treatments were randomly assigned to the aquaria to ensure balanced replication and minimize experimental bias.

### 2.2. Preparation of Papaya Seed Powder

Fresh seeds were manually extracted from ripe papaya (*Carica papaya* L.) fruits and thoroughly rinsed with clean water to remove any residual pulp or adhering membranes. The washed seeds were spread evenly on clean newspaper sheets and sun-dried for several days until a constant weight was achieved, indicating complete removal of surface moisture. The dried seeds were then pulverized into a fine powder using a laboratory electric grinder (Blender 800ES, Model BB90E). The obtained papaya seed powder (PSP) was packed in airtight polyethylene bags and stored in a cool, dry, and shaded environment until later use. The preparation method followed general protocols described for papaya seed processing and preservation [15]

### 2.3. Basal Feed Processing and Formulation

The basal diet for Nile tilapia (*Oreochromis niloticus*) was formulated using locally available ingredients, including fish meal, wheat bran, and barley bran. The raw fish used for meal preparation were first cooked for approximately one hour to eliminate pathogenic microorganisms, reduce excess oil content, and facilitate moisture removal. The cooked fish were then sun-dried for 3–4 days under hygienic conditions, ground into fine powder, and stored in airtight polyethylene bags until further use. The basal feed was formulated to contain 35% crude protein (CP), which is considered optimal for the growth performance and feed utilization of *O. niloticus*[16]. The crude protein contents of the individual ingredients were 50% for fish meal, 13% for wheat bran, and 10% for barley bran. [16, 17]. The average crude protein value of the bran mixture (wheat and barley) was therefore estimated at 11.5%. To achieve the desired 35% CP level, the proportions of each ingredient were calculated using the Pearson square method, a standard feed formulation technique widely employed in aquaculture nutrition.

The ingredients were weighed, thoroughly mixed with an appropriate amount of water to form a uniform dough, and sun-dried to a constant weight. The dried feed was subsequently ground into fine particles and stored in polyethylene bags for later use during the feeding trial. The general process of basal feed formulation from local ingredients is shown in Figure 2, while the composition and percentage contribution of each ingredient to the 35% crude protein diet are summarized in Table 1.

**Table 1.** Proportion (%) and Percent Contribution of Feed Ingredients to Crude Protein (35% CP)

| Feed Ingredient | Crude Protein (%) | Proportion in Diet (%) w/w) | Protein Contribution to Diet (%) CP) |
| --- | --- | --- | --- |
| Fish meal | 50.0 | 61 | 30.5 |
| Wheat bran | 13.0 | 19.48 | 2.53 |
| Barley bran | 10.0 | 19.48 | 1.95 |
| Total | — | 100% | 34.98 |

**Figure 2.**
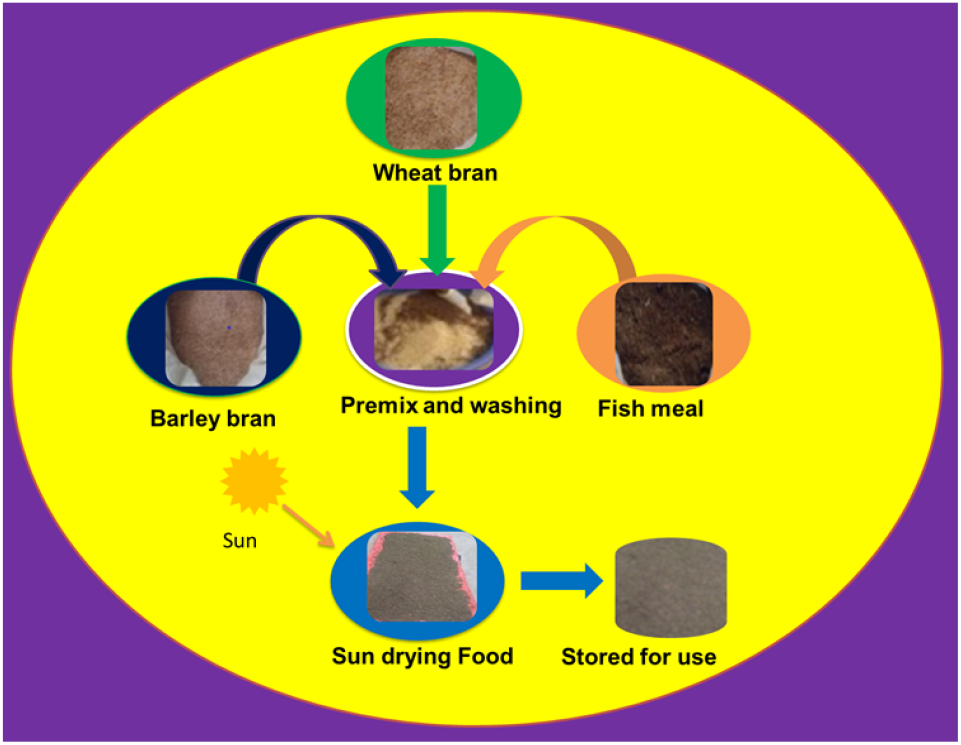
The process of feed formulation from local ingredients

### 2.4. Data Collection

#### 2.4.1. Growth Performance, Feed Utilization, and Survival Rate

Initial measurements of body weight, total length, and stocking number were recorded for fish in all eight experimental jars to evaluate growth performance, feed utilization efficiency, and survival rate. Measurements were taken every 15 days to adjust feed quantities and monitor growth trends throughout the experimental period [18, 19]. Total length (TL) was measured using a standard measuring board (to the nearest 0.1 cm), and body weight (W) was determined using an electronic balance (to the nearest 0.01 g). Growth performance was assessed using the Relative Growth Rate (RGR), which expresses the increase in body weight relative to the initial weight. Feed utilization efficiency was evaluated using the Feed Conversion Ratio (FCR), while survival rate (SR) was calculated based on the number of surviving fish at the end of the experiment[20].The following equations were used to calculate growth and survival indices [21]. These indices are widely recognized for assessing the biological performance and feed efficiency of fish under controlled aquaculture conditions.

Relative Growth Rate (RGR, %/day):

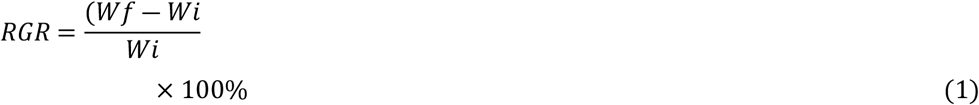

Where Wf= mean final weight and Wi= mean initial weight.

Feed Conversion Ratio (FCR):

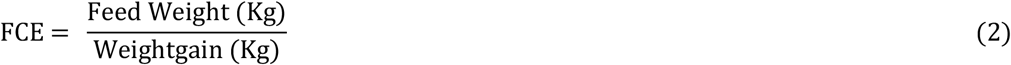

Survival Rate (SR, %) :

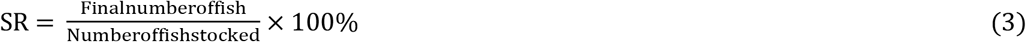

#### 2.4.2. Measurement of Reproductive Parameters

At the end of the 90-day experimental period, all fish from each tank were carefully collected using scoop nets and anesthetized in a diluted formalin solution prior to measurement. Each fish was assessed for total length (TL) and body weight (BW) using a measuring board and an electronic balance, respectively. After external measurements, the fish were dissected, and the gonads were carefully removed and weighed to determine gonad weight (GW).

The gonads were visually inspected for any morphological abnormalities or discolorations relative to the control group, following the standard morphological examination procedures described by [22]. Reproductive performance was evaluated using the Gonadosomatic Index (GSI), which reflects the proportion of gonad weight relative to total body weight and serves as an indicator of reproductive development and maturity [23]. The GSI was calculated using the following equation:

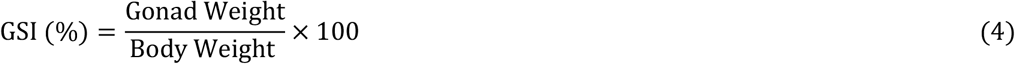

### 2.5. Physicochemical Parameters

Water quality parameters temperature, dissolved oxygen (DO), pH, and electrical conductivity (EC) were monitored weekly throughout the 90-days experimental period to ensure optimal rearing conditions for *Oreochromis niloticus*. Measurements were taken in each experimental tank using a portable multiparameter probe (HQd40, HACH, USA).

These parameters were maintained within the recommended ranges for tilapia culture as outlined by Boyd and Tucker (2012) and El-Sayed (2006). Continuous monitoring was essential to minimize environmental stress and ensure that any observed variations in fish growth and reproductive performance were attributable to dietary treatments rather than water quality fluctuations.

### 2.6. Statistical Analysis

Data collected for growth performance, feed utilization, survival rate, and reproductive parameters were statistically analyzed using one-way analysis of variance (ANOVA) with SPSS software (Version 23.0, IBM Corp., Armonk, NY, USA). Treatment means were compared using post hoc Tukey’s HSD tests to identify significant differences among groups. Statistical significance was accepted at p < 0.05, and all results were expressed as mean ± standard deviation (SD).

## 3. Results and Discussion

### 3.1. Growth Performance

The growth performance of *Oreochromis niloticus* fingerlings was significantly influenced by dietary supplementation with papaya seed powder (PSP) (Table 2). At the start of the experiment, the mean initial weights and lengths of fish did not differ significantly among treatments (P > 0.05), confirming uniformity in stocking size and eliminating initial size as a confounding factor. Initial weights in the Control, T1, T2, and T3 groups were 20.11 ± 4.77 g, 20.21 ± 4.82 g, 20.21 ± 4.82 g, and 20.67 ± 6.39 g, respectively, while the corresponding initial lengths were 10.04 ± 1.04 cm, 9.96 ± 0.81 cm, 9.96 ± 0.81 cm, and 10.52 ± 1.01 cm.

**Table 2.**
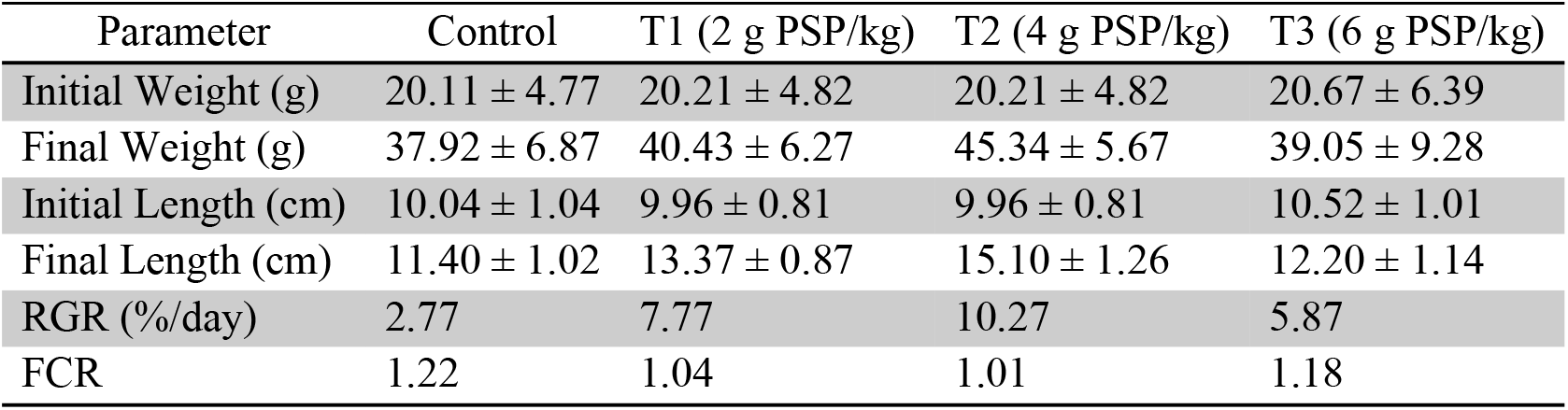
Growth performance, relative growth rate (RGR), and feed conversion ratio (FCR) of Nile tilapia fed diets supplemented with papaya seed powder (PSP).

After 90 days of feeding, significant differences (P < 0.05) were observed in final weights, lengths, and relative growth rates (RGR) among treatments. Fish fed the T2 diet (4 g PSP/kg) exhibited the highest growth, achieving a final weight of 45.34 ± 5.67 g, final length of 15.10 ± 1.26 cm, and RGR of 10.27% day^-1^. This was followed by T1 (2 g PSP/kg: 40.43 ± 6.27 g; 13.37 ± 0.87 cm; RGR = 7.77% day^-1^), T3 (6 g PSP/kg: 39.05 ± 9.28 g; 12.20 ± 1.14 cm; RGR = 5.87% day−^1^), and the Control group (37.92 ± 6.87 g; 11.40 ± 1.02 cm; RGR = 2.77% day^-1^). Feed conversion ratio (FCR) values ranged from 1.01 to 1.22, with the lowest FCR recorded in T2 (1.01), indicating the most efficient feed utilization. FCR values for all treatments fell within the optimal range (1.4– 1.8) reported for tilapia[24, 25], suggesting good diet quality and effective nutrient conversion.

The enhanced growth observed in the T2 group is likely due to the bioactive compounds present in PSP, including papain, proteolytic enzymes, flavonoids, and alkaloids, which may improve protein digestion, nutrient assimilation, and metabolic efficiency[15]. Papain, in particular, can hydrolyze proteins in the digestive tract, increasing the availability of amino acids for tissue growth, while flavonoids and other phytochemicals provide antioxidant support, reducing oxidative stress and promoting metabolic activity[26].

The dose-dependent response observed where growth decreased at higher PSP inclusion (T3, 6 g/kg), is likely due to anti nutritional factors such as tannins, saponins, and benzyl isothiocyanate, which can reduce feed palatability, inhibit digestive enzymes, and impairs nutrient absorption. Similar inhibitory effects have been reported in studies with plant-based feed additives, where moderate inclusion improved growth, but excessive amounts produced negative effects on metabolism and performance [26, 27].

The lower FCR in T2 indicates higher feed utilization efficiency, meaning that fish converted a larger proportion of feed into biomass. Efficient feed utilization is essential for aquaculture sustainability, reducing feed costs and optimizing production[28].

Additionally, PSP supplementation may modulate intestinal micro biota, promoting beneficial bacteria that enhance digestion and nutrient absorption[29]. This, together with enhanced proteolytic activity, likely contributed to the superior growth performance observed in the T2 group. Overall, these findings suggest that moderate supplementation of papaya seed powder at 4 g/kg diet can significantly enhance growth, feed efficiency, and nutrient utilization in Nile tilapia, while excessive inclusion may reduce these benefits due to anti nutritional effects.

### 3.2. Reproductive Parameters

The reproductive performance of *Oreochromis niloticus* was markedly influenced by dietary supplementation with papaya seed powder (PSP) (Table 3). In the control group, male and female gonad weights were 0.22 ± 0.16 g and 0.27 ± 0.25 g, respectively, with corresponding gonadosomatic index (GSI) values of 0.58 ± 0.38% for males and 0.84 ± 0.80% for females. In contrast, the treatment groups exhibited reduced gonad weights and GSI values. In T1 (2 g PSP/kg), male and female gonad weights were 0.04 ± 0.01 g and 0.18 ± 0.08 g, with GSI values of 0.08 ± 0.10% and 0.66 ± 0.02%, respectively. The T2 group (4 g PSP/kg) showed slightly lower gonad weights of 0.03 ± 0.02 g for males and 0.17 ± 0.21 g for females, with GSI values of 0.07 ± 0.10% and 0.43 ± 0.54%, respectively. In T3 (6 g PSP/kg), gonad weights were 0.08 ± 0.06 g for males and 0.19 ± 0.02 g for females, while GSI values were 0.19 ± 0.18% and 0.83 ± 0.28%, respectively.

**Table 3.**
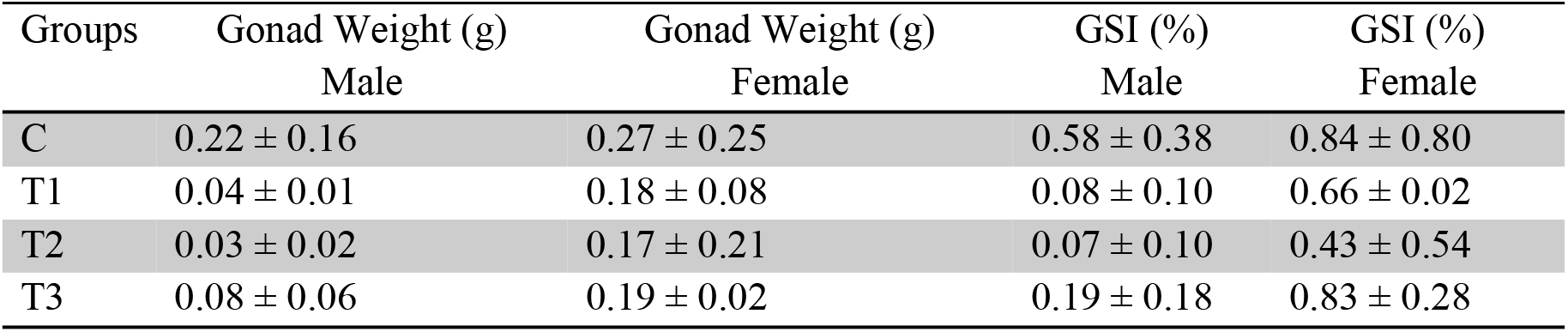
Mean ± SD reproductive parameters of Nile tilapia fed different inclusion levels of papaya seed powder (PSP) after 90 days.

The reductions in gonad weight and GSI across the treatment groups suggest a suppressive effect of PSP on gonadal development in both male and female tilapia. Compared to the control group, which recorded the highest gonad weights and GSI values, all PSP-supplemented diets led to a decline, indicating that increasing PSP levels may negatively affect reproductive performance.

These inhibitory effects are likely due to bioactive compounds present in papaya seeds, including benzyl isothiocyanate, papain, and certain alkaloids, which have been previously associated with anti fertility activity[15]. These compounds may interfere with steroidogenesis and gametogenesis, thereby reducing gonadal growth and function. The observed decrease in male GSI, particularly in T1 and T2, supports the potential role of PSP as a natural reproductive modulator in aquaculture species.

Similar effects have been reported in fish and mammals, where *Carica papaya* seed extracts were shown to inhibit gonadal development and reproductive hormone synthesis [26, 30]. The dose-dependent pattern observed in this study suggests that moderate PSP inclusion may suppress reproduction without causing overt toxicity, highlighting its potential application in managing breeding cycles in aquaculture. Therefore, the results indicate that while PSP can enhance growth and feed efficiency at moderate levels, its inclusion in the diet can adversely affect reproductive parameters, emphasizing the need to carefully balance nutritional benefits with potential reproductive suppression in Nile tilapia culture.

### 3.3. Water Quality Parameters

Throughout the 90-day feeding trial, physicochemical parameters of water were monitored to ensure optimal rearing conditions for *Oreochromis niloticus* (Table 4). Mean values for temperature, dissolved oxygen (DO), pH, and electrical conductivity (EC) remained within the acceptable range for tilapia culture. No significant differences (P > 0.05) were observed among the control and PSP-supplemented treatment groups, indicating that dietary inclusion of papaya seed powder did not adversely affect the aquatic environment.

**Table 4.** Mean ± SD physicochemical parameters of water in experimental tanks.

| Treatment | T ($^{\circ}\text{C}$) | DO (mg/L) | pH | EC ($\mu\text{S/cm}$) |
| --- | --- | --- | --- | --- |
| Control | $25.92 \pm 0.52$ | $5.42 \pm 1.37$ | $6.38 \pm 0.07$ | $130.80 \pm 19.75$ |
| T1 | $25.94 \pm 0.50$ | $5.08 \pm 1.53$ | $6.46 \pm 0.11$ | $134.98 \pm 18.63$ |
| T2 | 25.88 ± 0.50 | 5.74 ± 1.61 | 6.47 ± 0.09 | 129.69 ± 23.56 |
| T3 | 25.89 ± 0.48 | 5.97 ± 1.46 | 6.49 ± 0.09 | 129.62 ± 18.64 |

Water temperature ranged from 25.88 ± 0.50°C to 25.94 ± 0.50°C, within the optimal range for tilapia growth and metabolic activity. Dissolved oxygen levels varied between 5.08 ± 1.53 mg/L and 5.97 ± 1.46 mg/L, sufficient to support normal physiological functions and feeding behavior. The pH remained slightly acidic to near neutral, ranging from 6.38 ± 0.07 to 6.49 ± 0.09, which is acceptable for tilapia culture. Electrical conductivity values ranged from 129.62 ± 18.64 to 134.98 ± 18.63 µS/cm, suggesting stable ionic composition and minimal organic accumulation in all experimental tanks.

The stability of water quality parameters across all treatments indicates that variations observed in growth and reproductive performance were primarily attributable to dietary treatments rather than environmental factors. These findings are consistent with previous studies, such as Yadav et al. [10], who reported that inclusion of papaya seed in fish diets did not negatively impact water quality. Similarly, Farrag et al. [31] observed no significant differences in physicochemical parameters among treatment groups when plant-based feed additives were incorporated, suggesting that moderate inclusion of bioactive plant compounds does not compromise the aquatic environment.

Maintenance of optimal water quality is critical in aquaculture, as suboptimal conditions can impair metabolism, growth, and reproduction in fish. The results confirm that the experimental conditions provided a suitable environment for *O. niloticus*, allowing the observed effects on growth and reproduction to be attributed confidently to dietary PSP supplementation.

### 3.4. Survival Rate

The survival rate of *Oreochromis niloticus* fingerlings during the 90-day feeding trial was significantly influenced by dietary supplementation with papaya seed powder (PSP) (Table 5). Survival rates ranged from 65% in the control group to 90% in the T2 group. Specifically, the control group exhibited the lowest survival (65%), while T2 (4 g PSP/kg diet) recorded the highest survival (90%). Survival rates in T1 (2 g PSP/kg) and T3 (6 g PSP/kg) were 80% and 75%, respectively, indicating that moderate PSP supplementation enhanced fish viability, whereas excessively high inclusion (T3) slightly reduced the benefit.

**Table 5.** Survival rate of Nile tilapia fingerlings fed diets supplemented with papaya seed powder (PSP).

| Treatment | Number of Stocked Fish | Number of Harvested Fish | Survival Rate (%) |
| --- | --- | --- | --- |
| Control | 20 | 13 | 65 |
| T1 | 20 | 15 | 80 |
| T2 | 20 | 18 | 90 |
| T3 | 20 | 16 | 75 |

The increased survival rates in T1, T2, and T3 compared to the control suggest that dietary inclusion of PSP confers protective effects on fish health. The highest survival in T2 indicates an optimal supplementation level, whereas the slight reduction in T3 may be due to saturation effects or minor anti-nutritional factors, such as tannins and saponins, present in papaya seeds at higher inclusion rates.

These findings are consistent with previous reports indicating that papaya seeds contain bioactive compounds with antimicrobial, antioxidant, and immune-stimulatory properties, which enhance fish resilience to environmental stressors and pathogenic challenges[15, 31]. Such effects likely contributed to the improved survival observed in T1 and T2. The results highlight the potential of PSP as a natural feed additive to improve health, reduce mortality, and promote sustainable aquaculture practices.

## 4. Conclusion and future recommendation

This study demonstrated that dietary supplementation with papaya seed powder (PSP) significantly influenced the growth, survival, and reproductive performance of *Oreochromis niloticus*. The optimal growth performance and highest survival rate (90%) were observed at a moderate inclusion level of 4 g PSP/kg feed, indicating that this concentration enhances feed utilization, nutrient assimilation, and overall fish health.

However, higher inclusion levels of PSP resulted in reduced growth and gonadal development, suggesting a threshold beyond which papaya seed powder may exert adverse effects. The observed decline in gonad weight and gonadosomatic index (GSI) in PSP-fed groups highlights its potential as a natural reproductive inhibitor, which could be strategically applied for population control in tilapia aquaculture. Therefore, papaya seed powder appears to be a cost-effective and eco-friendly feed additive that promotes growth and survival at optimal levels while modulating reproductive performance. Based on these findings, PSP can be considered a promising natural agent for enhancing growth and managing reproduction in *O. niloticus*, particularly in aquaculture systems where population management is essential. Further study is needed to assess the well-being of the experimental fish, as certain parasites were observed in the current research. Additionally, a histopathological study is required to determine any influence of the bio-active material in PSP on the tissues of the experimental fish.

## Acknowledgment

We kindly acknowledge Department of Biology, Mattu University, for providing the space for the experiment.

